# Exploring the impact of inoculum dose on host immunity and morbidity to inform model-based vaccine design

**DOI:** 10.1101/328559

**Authors:** Andreas Handel, Yan Li, Brian McKay, Kasia A. Pawelek, Veronika Zarnitsyna, Rustom Antia

## Abstract

**Background:** Vaccination is an effective method to protect against infectious diseases. An important consideration in any vaccine formulation is the inoculum dose, i.e., amount of antigen or live attenuated pathogen that is used. Higher levels generally lead to better stimulation of the immune response but might cause more severe side effects and allow for less population coverage in the presence of vaccine shortages. Determining the optimal amount of inoculum dose is an important component of rational vaccine design. A combination of mathematical models with experimental data can help determine the impact of the inoculum dose.

**Methods:** We designed mathematical models and fit them to data from influenza A virus (IAV) infection of mice and human parainfluenza virus (HPIV) of cotton rats at different inoculum doses. We used the model to predict the level of immune protection and morbidity for different inoculum doses and to explore what an optimal inoculum dose might be.

**Results:** We show how a framework that combines mathematical models with experimental data can be used to study the impact of inoculum dose on important outcomes such as immune protection and morbidity. We find that the impact of inoculum dose on immune protection and morbidity depends on the pathogen and both protection and morbidity do not always increase with increasing inoculum dose. An intermediate inoculum dose can provide the best balance between immune protection and morbidity, though this depends on the specific weighting of protection and morbidity.

**Conclusions:** Once vaccine design goals are specified with required levels of protection and acceptable levels of morbidity, our proposed framework which combines data and models can help in the rational design of vaccines and determination of the optimal amount of inoculum.

## Introduction

Vaccines are the best and most cost-effective defenses we have against many infectious diseases. While the composition of a vaccine can be complex, the most important component is the antigen of the pathogen against which one wants to immunize [1]. Different types of vaccines exist, those based on antigens that contain the pathogen in a non-replicating form, and those that contain the pathogen in a replicating form, usually attenuated to reduce morbidity and mortality [1].

When deciding on the inoculum dose for a vaccine, one often needs to strike a balance between conflicting goals. Higher doses generally lead to more immunity and better protection [2]. Lower doses might reduce vaccine side effects and might also be required if there is a vaccine shortage, for instance due to a pandemic emergency, manufacturing issues or high costs [3,4]. The ability to predict how changes in inoculum dose impact immune protection and morbidity, and how to achieve the best balance between enough inoculum to trigger a robust immune response and low enough inoculum would significantly contribute toward better vaccine design [5–12].

Currently, the main way to determine vaccine inoculum dose is by trial and error, which is expensive and logistically challenging [13–16]. A way to improve this approach is to combine mathematical models with experimental data. Such approaches are commonly applied to drugs, where pharmacokinetic/pharmacodynamic (PK/PD) models are used in combination with experimental data to try and optimize drug dosing regimens [17]. Application of a similar approach to vaccines has been recently proposed for tuberculosis [18].

Here, we develop and analyze a quantitative modeling framework that might allow us to eventually predict the optimal inoculum dose for a given vaccine and setting. We develop our modeling framework for live attenuated vaccines using data from two infection experiments, namely influenza A virus (IAV) and human parainfluenza virus (HPIV). We further investigate a scenario for an inactivated vaccine.

Influenza A virus remains a serious health concern. While a vaccine exists, it needs to be reformulated regularly. Even when the vaccine is well-matched to the circulating strain, its efficacy is not as good as that of other vaccines, especially in the elderly. It has been suggested that using a higher inoculum dose in vaccines for this population might be beneficial [19]. Development of a better vaccine that remains protective in the presence of antigenic drift and that has a higher efficacy remains a priority.

Human parainfluenza virus (HPIV) is an important cause of lower respiratory tract illness in children [20–24]. There is currently no licensed vaccine available against HPIV [11,21,24], despite various attempts to develop such a vaccine [25].

While the two pathogens we analyze here are important on their own, we consider the most important contribution of this study to be the development of a conceptual, quantitative framework that may be used to rationally design vaccines and determine an optimal inoculum dose for any pathogen.

## Materials and Methods

### Experimental data

We analyzed data from two previously published studies, one on influenza A virus (IAV) infections in mice [26] and the other on human parainfluenza virus (HPIV) type 3 infection in cotton rats [27].

For the IAV study, groups of mice were infected with 6 different inoculum doses of the H1N1 PR8 strain of influenza. Geometric mean viral titers were recorded at different times following the infection with each dose. In addition, lung damage was measured and scored.

For the HPIV study, groups of cotton rats were infected with 5 different doses of HPIV-3. Geometric mean viral titers were recorded at different times following infection in both lung and nose. For the highest inoculum dose, the study additionally reported several virus measurements over the first 96 hours. The study also reported antibody titers 21 days after infection for the 3 lowest inoculum doses for which virus data was reported.

We used an additional data set to estimate a mapping between innate immune response strength and morbidity. This data was taken from a previously reported challenge study of influenza infection in human volunteers [28]. We used the reported values for different components of the innate response (IFN-a, IL6, IL8 and TNF-a) and total symptom score as measure of morbidity.

For further experimental details, we refer the reader to the original studies.

### Mechanistic dynamical infection model

We formulated and implemented a mechanistic, dynamical model of the infection dynamics based on a set of ordinary differential equations. The model is based on our previous work, where we analyzed the relationship between inoculum dose and viral load dynamics [29]. The model is also similar to many other models that have recently been used to model acute viral infections (see e.g. [30–33]).

Our model tracks target cells, virus, and certain immune response components. Uninfected cells, *U*, become infected by free virus, *V*, at rate *b*. Infected cells, /, produce virus at rate *p* and die at rate *d_I_*. For purposes of comparison with the data, we keep track of dead cells through an extra compartment, *D*.

Free virus infects cells at rate *b′*, is cleared by antibodies at rate 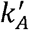 or removed due to other mechanisms (e.g. mechanical transport) at rate *d_v_*. Note that *b′* and 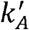 differ from parameters *b* and *k_A_* to account for experimental units (PFU for virus and titer for antibody). Since we are modeling short, acute infections, we follow the usual assumption and ignore growth and death of uninfected target cells [30,31].

In addition to the basic infection process, we also model components of the innate and adaptive immune response. We consider a generic innate response, *F*, which is produced and decays at rates *p_F_* and *d_F_* in the absence of an infection. Presence of virus leads to an increase in the innate response, with growth saturating at a maximum rate *g_F_*. The maximum level the innate response can reach is given by the saturation parameter *F*_max_. Since the innate response units are arbitrary, the model is set up such that in the absence of infection, the innate response is at a steady level of *F* = 1, which leads to *p_F_* = *d_F_*. We also fix the parameter representing the decay rate at *d_F_* = 1 per day, which is in line with estimates from an influenza infection analysis in ponies [34].

The innate response is modeled as having two main mechanisms of action. First, it can directly counteract the virus by, for instance, reducing virus production rate of infected cells [35]. In our model, the strength of production suppression is determined by the parameter *s_F_*. The second action of the innate response is to induce the adaptive response, as described next.

For the adaptive response, we focus on B-cells and antibodies, which are the major correlates of protective immunity for most vaccines, including HPIV and IAV [23,36]. The dynamics of activated B cells is modeled as increasing in a sigmoidal manner dependent on both the amount of virus (antigen) and the innate response, with a maximum rate *g_B_*. Since we are focusing on the short-term dynamics of the system, B-cell decay is ignored. In the absence of an infection, B-cells are set to an arbitrary level of 1. B-cells produce antibodies at rate *r_A_*. Antibodies decay naturally at rate *d_A_* and bind to and remove free virus at rate *k_A_*.

The model is implemented as a set of ordinary differential equations given by the following set of equations:

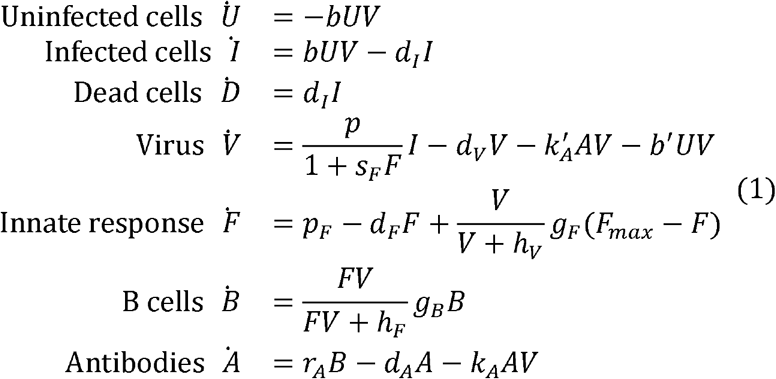

### Model fitting

The model is fit to the IAV and HPIV data. For IAV the fit is to the virus load and lung damage data. For HPIV, the fit is to the virus load and antibody data. For each pathogen, we fit data for all different inoculum doses simultaneously to the model. For each inoculum dose, *i*, we estimate the starting value for the virus inoculum, *V_i_*. All other model parameters are shared across different inoculum doses.

Model performance is assessed by the sum of squared residuals (SSR). To allow computation of a single SSR value for the different experimental variables, the contribution of each variable is non-dimensionalized by dividing by the variance of the data. To give the different experimental variables comparable importance, we also divide each variable by the number of data points. This amounts to over-weighting the few data points for lung damage (AIV) and antibody response (HPIV) and reducing the weight for the more plentiful viral load data. Mathematically, the expression for the SSR is given by

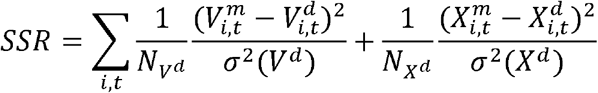

Here, *V* is viral load (on a log scale) and *X* represents either antibodies (for HPIV) or damage (for IAV), the superscript indicates model (*m*) or data (*d*), the sum runs over all inoculum doses, *i*, and all time points, *t*. *N* indicates the number of data points for either the virus or the other variable, *σ^2^* indicates the variance for that variable. Since both damage and antibodies (*X*) are measured in different units in the data and the model, each are normalized before subtracting and squaring. While this re-scaling is only necessary for the instances where we compare model and date (lung damage for IAV and antibodies for HPIV), for consistency between models, we show re-scaled values for both lung damage and antibodies for both AIV and HPIV.

The model is being fit by varying model parameters to minimize the *SSR*. When doing so, we take into account left-censored nature of the data. If the reported virus load is at or below the limit of detection (LOD, which is 0.27 loglO units for IAV and 2 loglO units for HPIV as reported in the original studies), we treat the difference between model and data as being the difference between model and LOD if the model prediction is above the data, and we do not count any difference between model and data for any model prediction below the LOD data point [37,38].

### Model implementation

All computations were done in the R programming language version 3.4.3 [39]. Fitting was done using the nloptr optimizer package [40], differential equations were integrated using the deSolve package [41]. All data and code required to reproduce all results presented here are supplied as supplementary material.

## Results

### Data extraction

The data used for our study was obtained from the original reports as follows.

For the IAV study, we obtained log viral load and lung lesion score expressed in percent lung damage from table 1 of [26]. The viral kinetics of the highest inoculum dose strongly hints at survivor bias (see figure 1 of [26]). Specifically, the data suggest that sicker mice, with presumably higher virus load, were killed and sampled first, while less sick mice, with presumably lower virus load, were kept alive and sampled later. We, therefore, decided to exclude the data for the highest inoculum dose from consideration, leaving us with viral load and percent lung damage data for 5 different inoculum doses.

**Figure 1.**
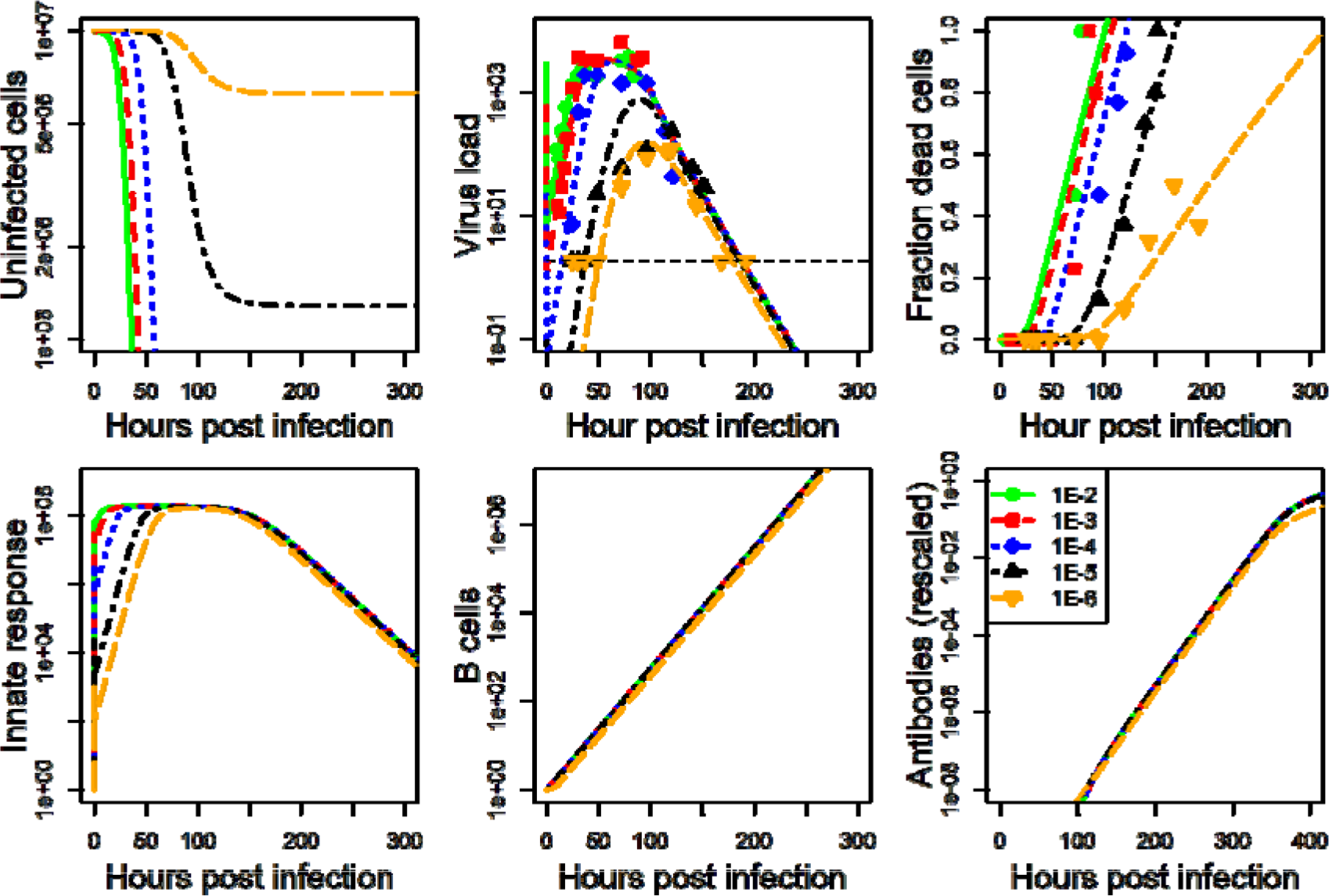
IAV infection at five different inoculum doses. Data was available for virus load and cell damage. Kinetics for 6 of the seven model compartments for the best fit model are shown. Infected cells kinetics very closely follows virus kinetics and is therefore not shown. Dashed horizontal line indicates the limit of detection for virus load.

For the HPIV study, we focused on lung viral load. The data was extracted from figures 1 and 2 of [27] using Engauge Digitizer [42]. Viral load kinetics for the highest inoculum were measured twice with some overlap in times (24h and 96h). We averaged data for these times from the 2 experiments. We additionally obtained data on antibody titers for those inoculum doses for which viral load was reported (the 3 lowest inoculum doses) from figure 3 of [27].

**Figure 2.**
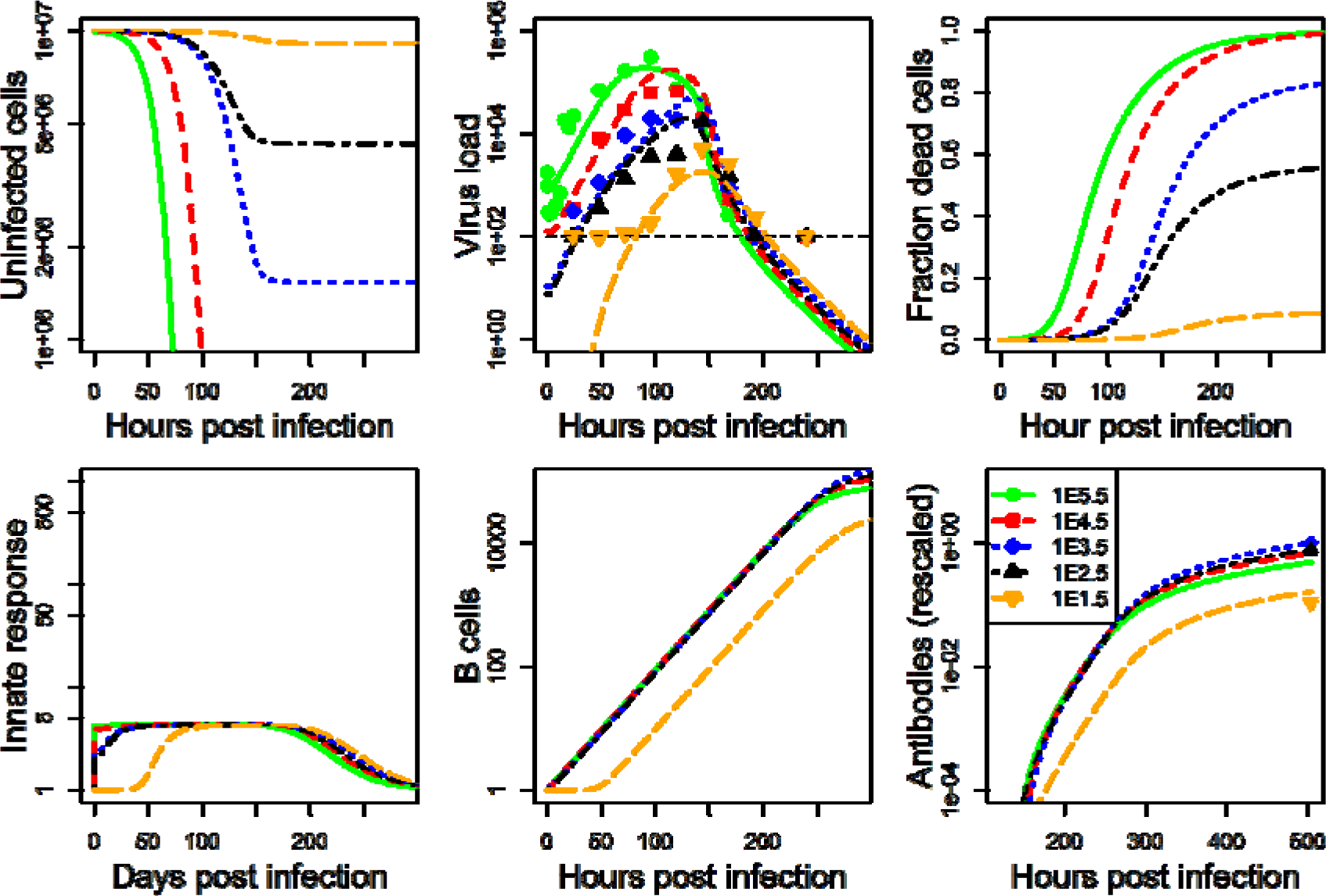
HPIV infection at five different inoculum doses. Data was available for virus load and antibody titers. Kinetics for 6 of the seven model compartments for the best fit model are shown. Infected cells kinetics very closely follow virus kinetics and is therefore not shown. Dashed horizontal line indicates the limit of detection for virus load.

**Figure 3.**
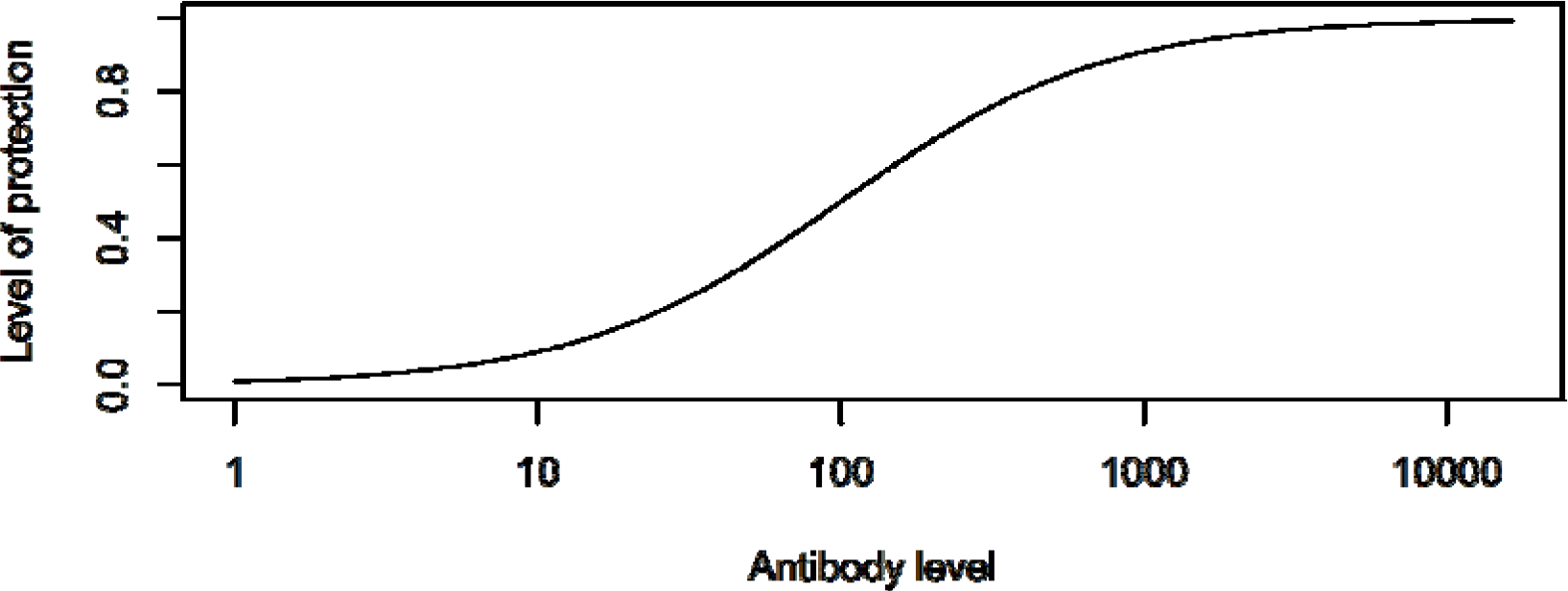
Protection as function of antibody levels (k_1_ = 1, k_2_ = log(100)).

For the data linking innate response to symptoms, total symptoms score data was extracted from figure 3 and innate immune response data from figures 5 and 6 of [28] using Engauge Digitizer. More details on how this data was used are provided in a later section.

Data extracted from these 3 studies are shown in Figures 1, 2 and 4 (together with the best fit models, described below), and are also included in the supplementary material.

### Model development and fitting

The model we developed is described in detail in the methods section. For each data set, we fit the model to the viral load data, and either lung damage (IAV) or antibody (HPIV) data. Details on the fitting approach are provided in the methods section. The model fits and data for IAV and HPIV are shown in figures 1 and 2 respectively. Parameter values for the best fits are given in the supplementary material.

### Quantifying Immune Protection

We want to quantify the amount of protective immunity induced by different inoculum doses. We focus on the B-cell and antibody component of the adaptive immune response. Provided antibodies are specific to the pathogen, higher levels of antibodies generally lead to better protection [43–45], Recent studies for influenza vaccines [46,47] have shown that the following function provides a good mapping from antibody titer to the level of protection from infection:

Here, the level of protection, , varies between 0 and 1, with low protection for low levels of antibody titer, , and maximum protection at high levels. The constants and determine the slope of the curve and the level at which protection is at 50% respectively (see [46] for more details). This functional shape is also consistent with data for other pathogens [43–45], Figure 3 illustrates this relationship between antibody levels and protection graphically.

Our model represents antibodies in units of numbers of antibodies. In general, experimental studies report antibody neutralizing titers or similar assay-specific units. For this reason, and because we have no data for the correlation between antibody titers and protection for either the HPIV or IAV data we analyze, it is impossible to determine specific choices for and for our study systems. We instead chose values such that the antibody levels considered span the full range from low to high protection levels. Specifically, we set and where is the range of antibody levels predicted by our model for different inoculum doses and is the expected value. This choice is essentially arbitrary and therefore the protection curves we present below are to be understood conceptually.

### Quantifying Morbidity

It is still not fully understood how virus and immune response affect host morbidity, i.e. the severity of symptoms. For virus infections, host morbidity can result in virus-induced death of infected cells, as well as immune response mediated pathology. A study of influenza infection in humans showed that a model in which symptom score was proportional to innate cytokine levels provided an adequate fit to the data [48], Another study of influenza infections used a combination of innate cytokine (interferon) levels and cell death to define morbidity as, where is the total number of dead cells, and was chosen to be a sigmoidal mapping of log interferon levels [49], Similarly, a previous model for dengue infections assumed that morbidity was proportional to the peak of the innate response, i.e. [50], In the case of vaccines, strong pathological effects such as the death of a meaningful fraction of target cells do not occur. It therefore seems most reasonable to express morbidity (strength of symptoms) as a function of the innate immune response.

To obtain an estimate for a mapping between innate immune response and morbidity, we use data from a previously reported challenge experiment of influenza infection in human volunteers [28], We use the reported values for different components of the local innate response (IFN-a, IL6, IL8, and TNF-a) and, after scaling each component to a maximum value of 1, sum them to obtain an estimate for the total innate response strength. This total response quantity is again scaled and then mapped to morbidity, measured as total symptom score. Figure 4 shows the data.

**Figure 4.**
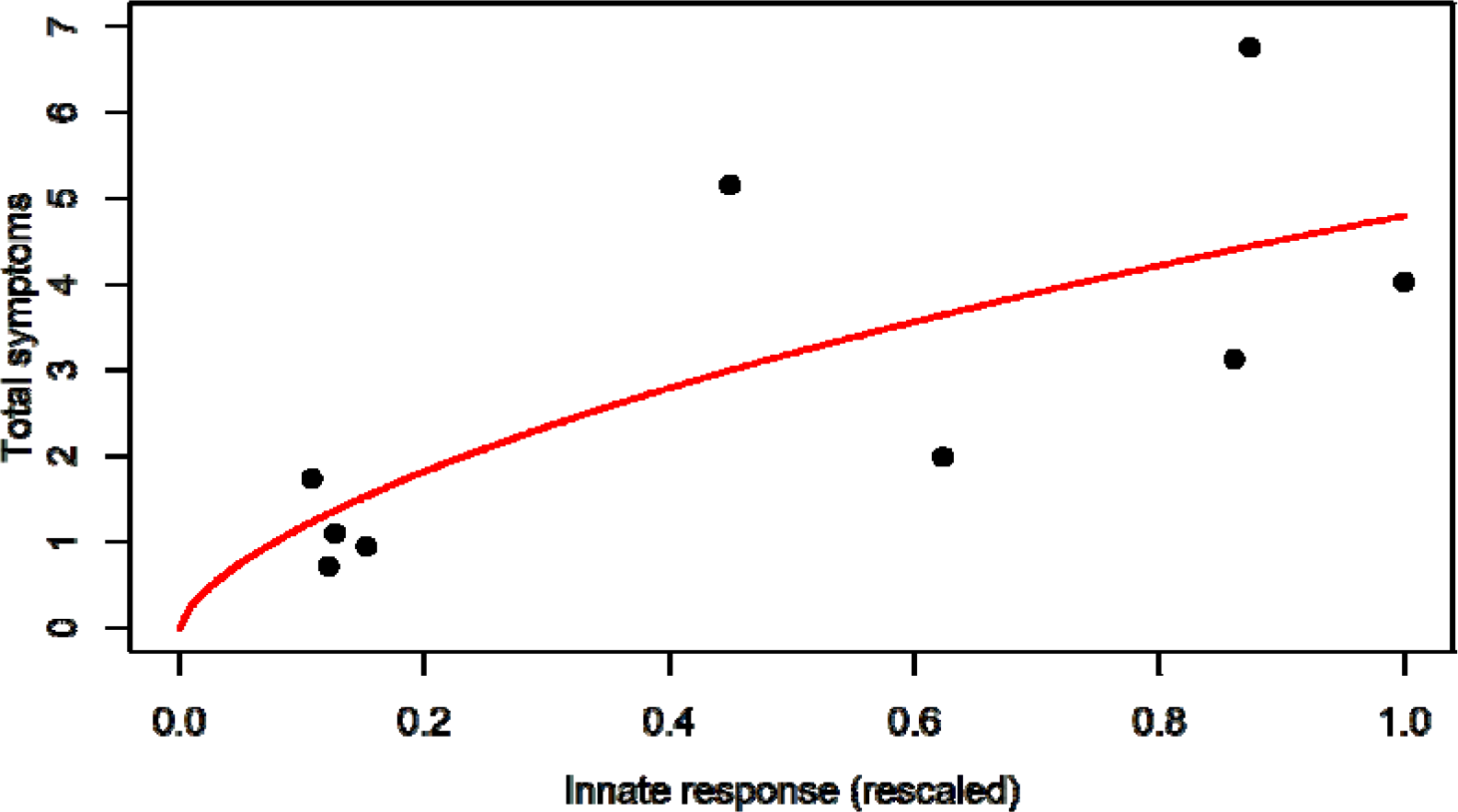
Data and best fit model for the connection between immune response and symptoms.

Also shown is the best fit of a sigmoidal model that provides a mapping between innate response and morbidity. The model is given by where is morbidity as measured by total symptom score, and is the scaled innate response. Best fit parameter values are = 6.5 and = 0.66. The parameter was fixed at 36, corresponding to the maximum score possible based on the study protocol [28], While a simpler linear model would fit the data equally well, it is less biologically reasonable since it would allow an unbounded increase in symptoms.

From our simulations, we obtain the time course of the innate response,. After rescaling this quantity, we use equation (3) to compute the time course for morbidity,. Finally, we take the integral of the morbidity over the duration of the infection to compute total morbidity as the area under the morbidity curve (MAUC). Since this approach mixes model simulations based on animal infections with morbidity estimates based on human data, the resulting morbidity curve should be interpreted in a similar conceptual way as the protection curve described above.

### Immunity and pathogenesis as function of inoculum

After fitting the dynamical infection model (1) to each data set, we used the best-fit parameter values and ran simulations for a range of inoculum doses. Several time-series for the IAV and HPIV model simulations spanning the whole range of simulated inoculum doses are shown in figures 5 and 6.

**Figure 5.**
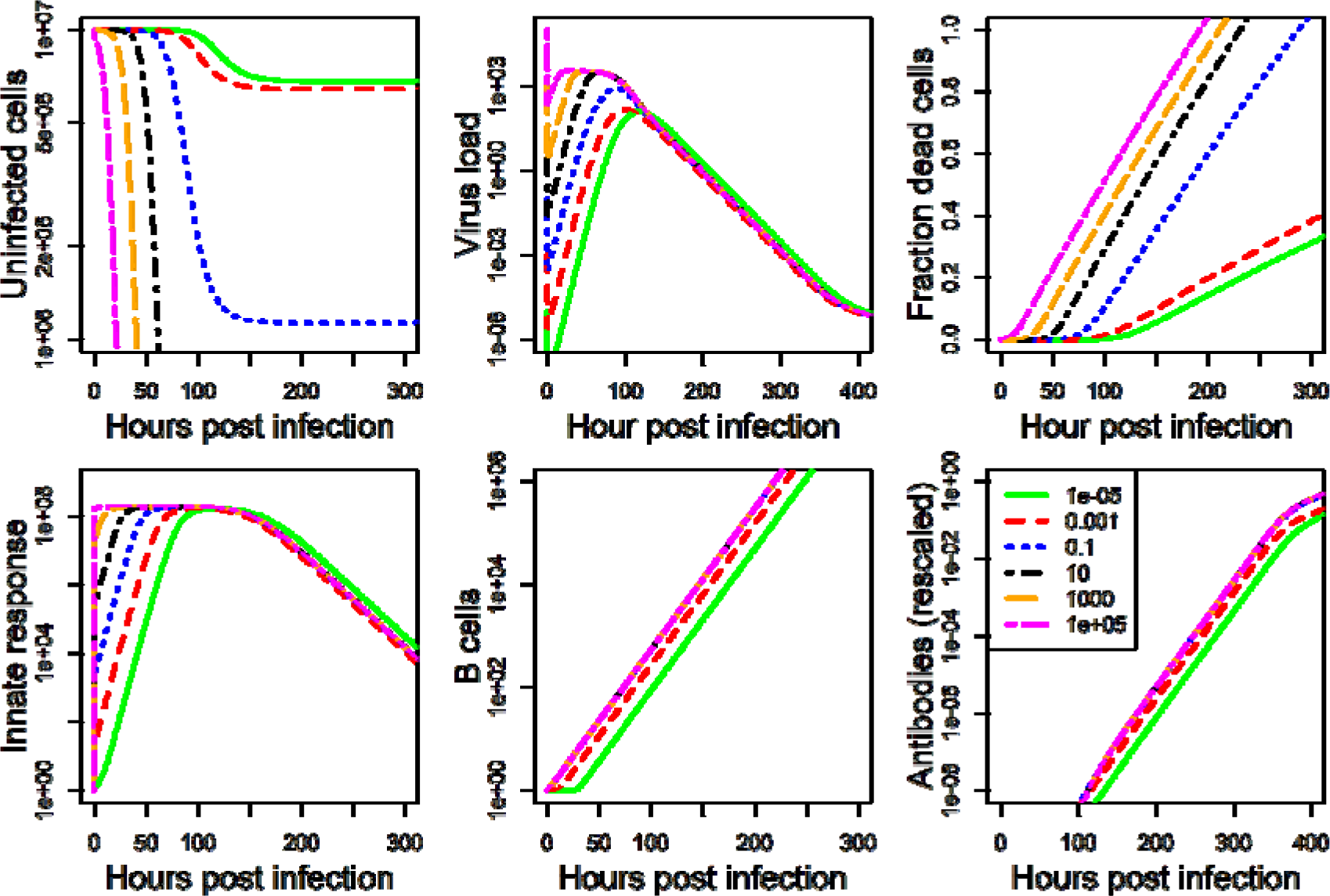
IAV model simulation for a range of inoculum doses.

**Figure 6.**
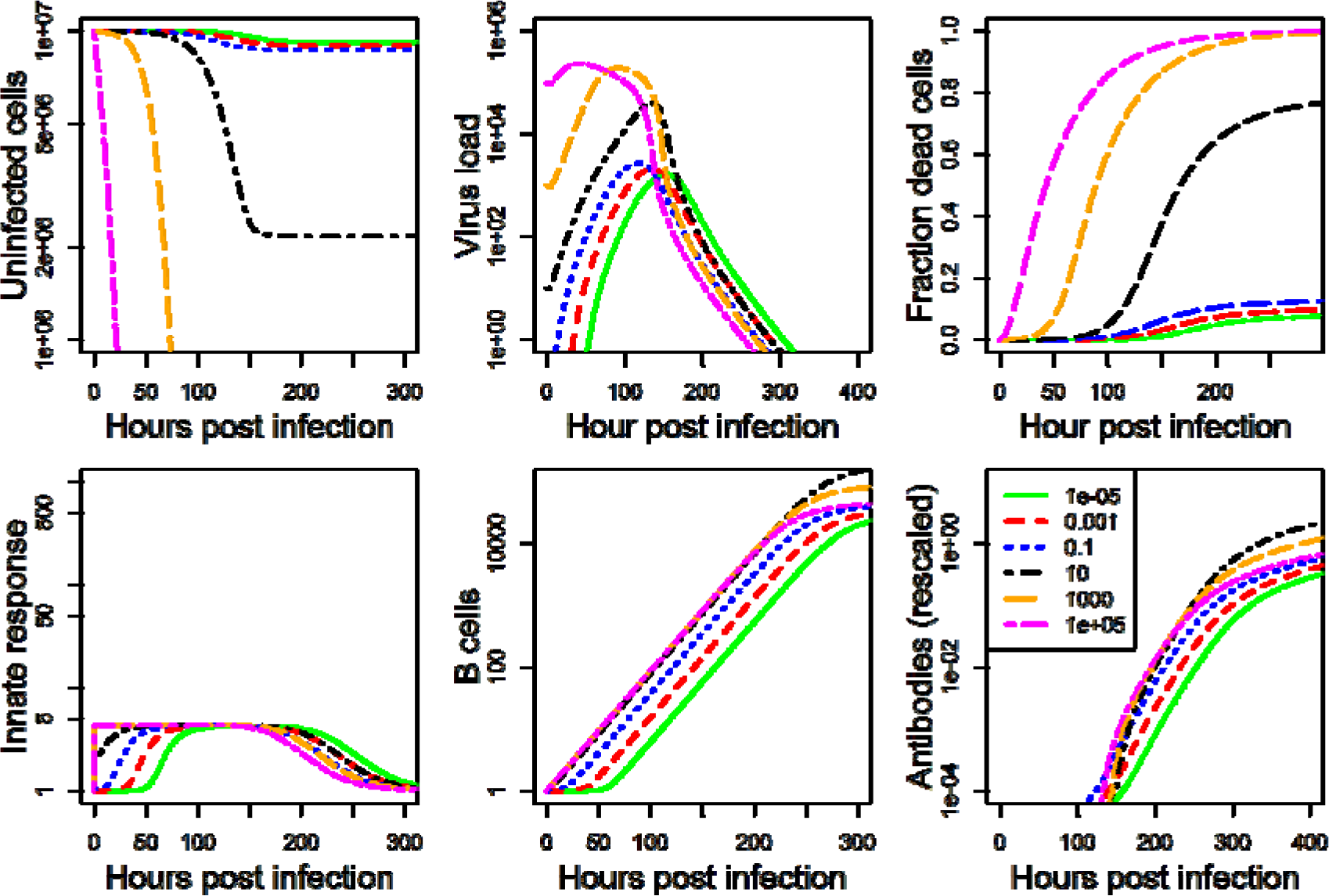
HPIV model simulation for a range of inoculum doses.

From these time-series, the level of protection and morbidity was computed. The model is simulated for 21 days, predicted antibodies are recorded at the final time. From these antibody levels, we compute immune protection using equation (2). We also record the predicted innate immune response, and, after scaling, use equation (3) to compute morbidity, and by integrating the area under the curve, determine the total amount of morbidity during the infection. Those results are shown for IAV are shown in figure 7, figure 8 shows results for HPIV.

**Figure 7.**
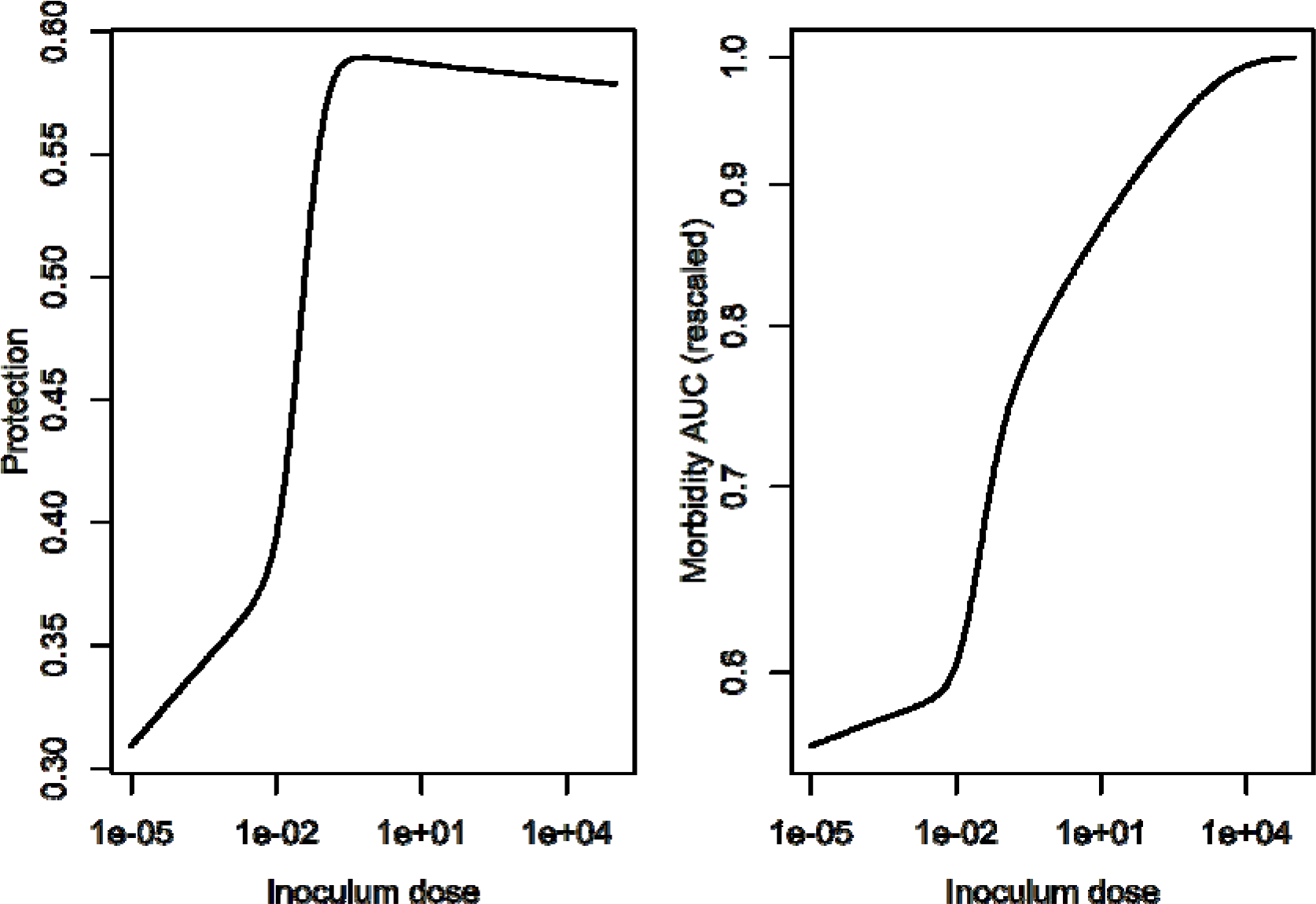
Inoculum dependent protection and damage for the IAV infection model.

**Figure 8.**
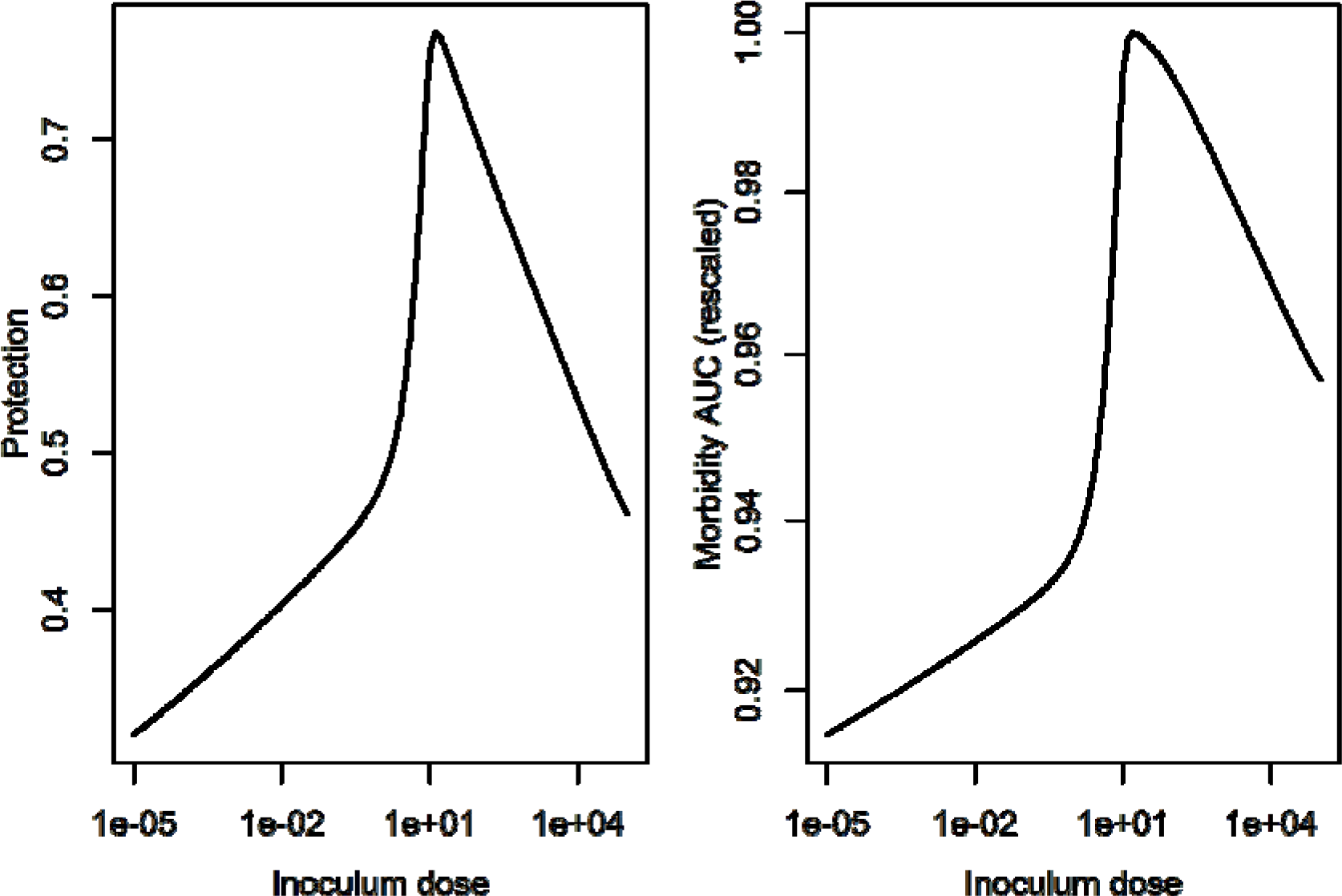
Inoculum dependent protection and damage for the HPIV infection model.

### An inactivated vaccine model

The model and data above are for replicating pathogens, as such representing live, attenuated vaccines. Another important category of vaccines are those where the pathogen is killed and non-replicating. A modification of the above model can be used to simulate such a vaccine. For such a vaccine, cells do not get productively infected, and one can remove the variables tracking uninfected and infected cells. The model simplifies to

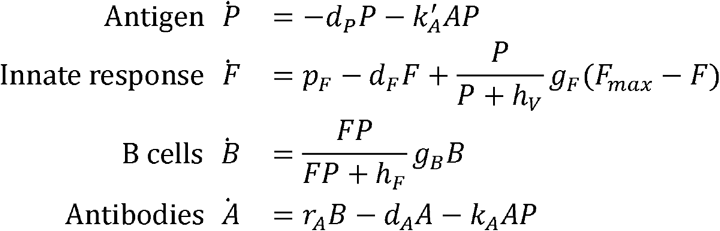

We were not able to find data in the published literature for antigen (and possibly other model components) time series for different inoculum doses that would be detailed enough to allow model fitting. We therefore instead chose arbitrary values for model parameters that produced reasonable dynamics and explored the impact of inoculum/antigen dose on protection and morbidity for such a generic model. Figure 9 shows simulated time-series for different inoculum doses and figure 10 shows the resulting predicted immune protection and morbidity.

**Figure 9.**
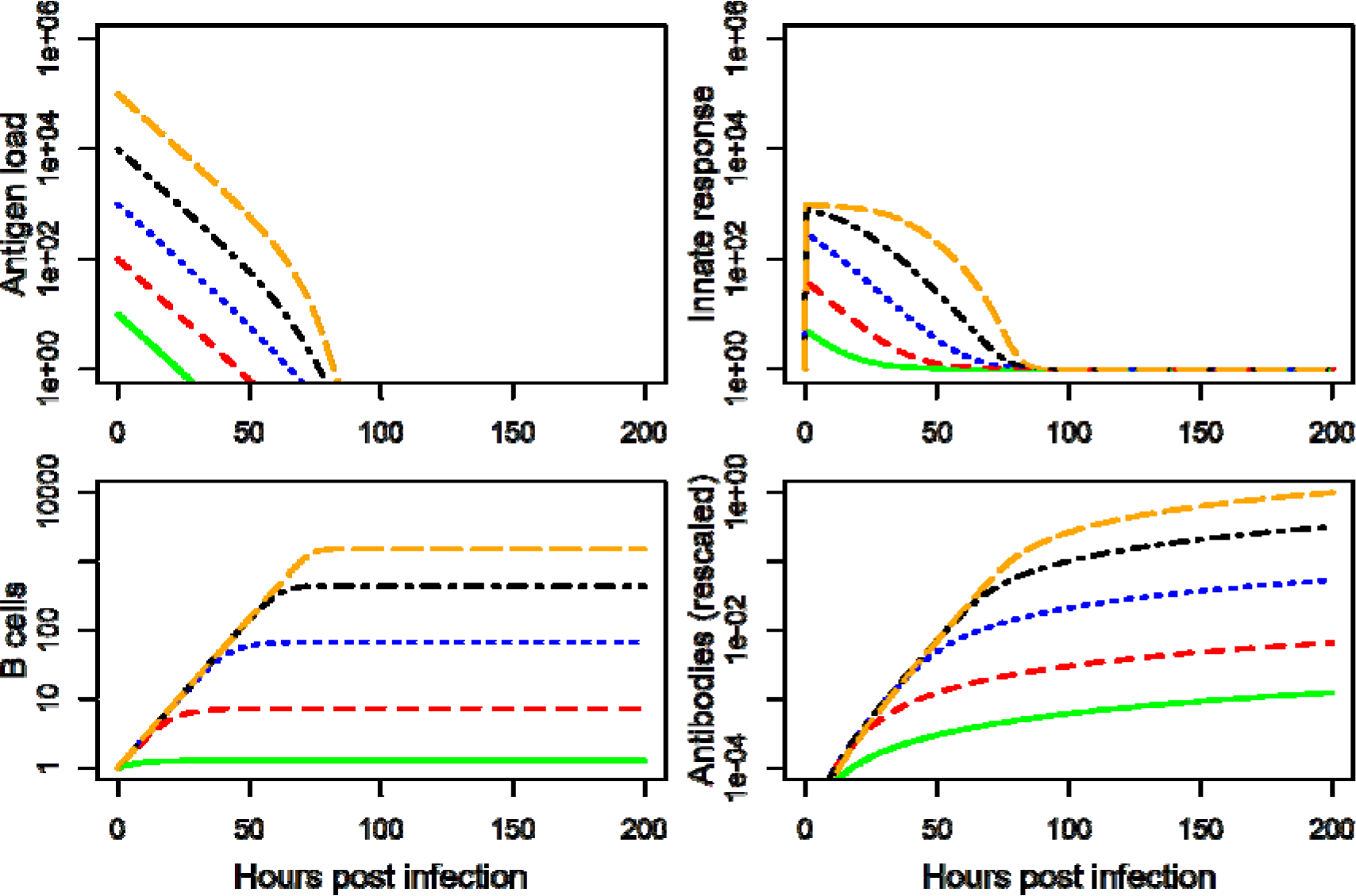
Model for non-replicating vaccine. Model parameters were set to. Initial conditions are F = 1, B = 1, A = 0 and varying values for antigen load.

**Figure 10.**
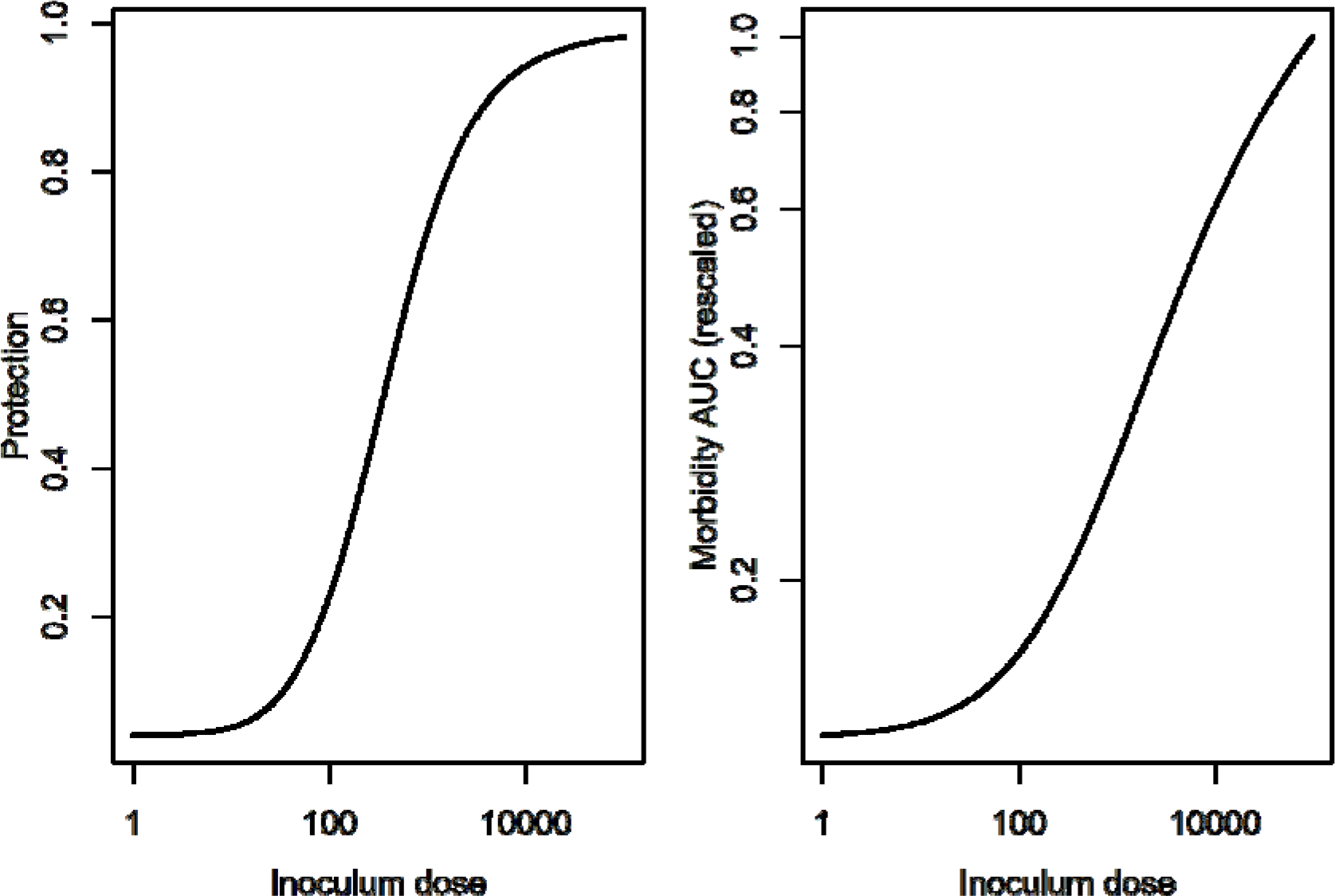
Inoculum dependent protection and damage for the inactivated vaccine model.

### Optimal Inoculum dose illustration

Once immune protection and morbidity as a function of inoculum dose are predicted by the model, one can potentially determine optimal inoculum dose choices. The optimal amount depends on the main goals of the vaccine formulation. One could, for instance, choose a minimum acceptable level of immune protection or maximum acceptable level of morbidity, and determine the inoculum dose for those criteria. Another possibility is to compute and maximize a quantity that is a compound of immune protection and morbidity, with specific weights assigned to protection and morbidity. We illustrate this idea conceptually by looking at a very simple quantity, namely the ratio of immune protection to morbidity (as defined by the area under the curve), P/M. Figure 11 shows this quantity for the IAV and HPIV infections as well as for the inactivated vaccine.

**Figure 11.**
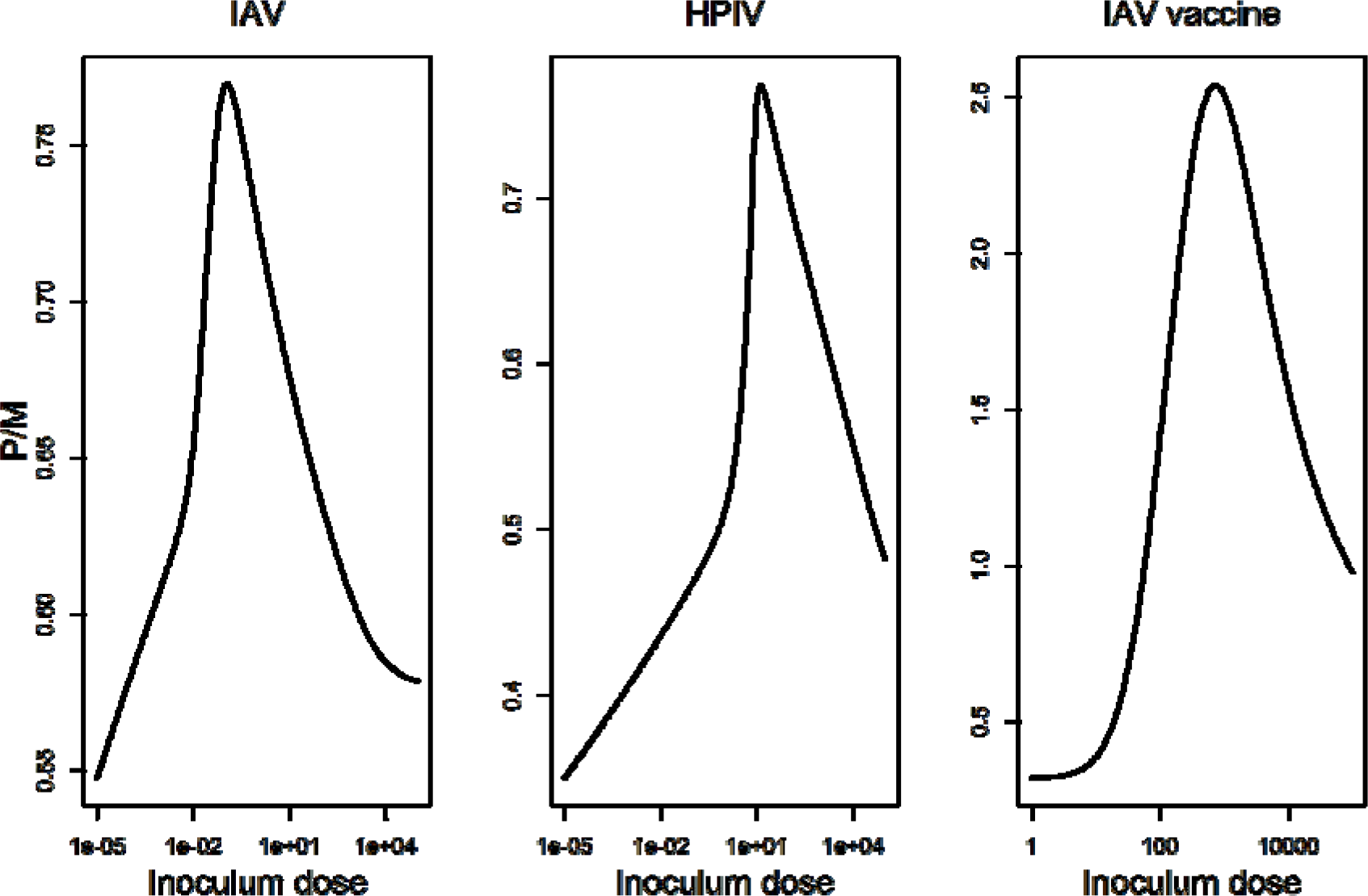
Ratio of protection, P, over morbidity, M, for different inoculum doses.

In each case, the amount of inoculum that leads to the highest ratio of P/MAUC occurs at an intermediate dose.

## Discussion

In many situations, a higher inoculum dose of either live attenuated or killed antigen in a vaccine leads to a stronger immune response and subsequently likely better antibody or T-cell mediated immune protection [51]. However, this does not have to be universally true. Once inoculum doses increase beyond some threshold, the innate immune response might be triggered too strongly, which in turn could lead to an impaired adaptive immune response and thus reduced immune protection. Similarly, while increased inoculum usually leads to more morbidity and stronger symptoms, this again might not be universal and depends on the interaction of pathogen and immune response.

In this study, we used a combination of data and models to explore how inoculum dose impacts immune protection and morbidity. We found that for the examples investigated, sometimes there is a monotonic or almost monotonic increase of protection and morbidity as inoculum increases (inactivated vaccine and IAV examples), while other times protection and morbidity can decline once dose increases beyond some value (HPIV example). For the illustrative example of an optimal dose based on a simple ratio of protection to morbidity, all our examples suggest that an intermediate amount of inoculum dose is optimal.

Our study fits into the recently proposed framework of Immunostimulation/Immunodynamic (IS/ID) modelling, which has been proposed as a framework to combine models and data for better vaccine formulation decisions [18], in analogy to the well-established pharmacokinetic/pharmacodynamic (PK/PD) modelling approach widely used in drug development [17].

We believe that using an approach that combines modeling with data can help in the development of more efficient vaccines. The key toward that goal is the availability and integration of the right kind of models and data. Since the data we analyzed is a mix of animal and human data and neither is complete enough to allow the whole modeling framework to be applied to it, the results presented here are to be considered mainly conceptually. The ideal type of data tracks antigen or pathogen load and different immune components over time, as well as morbidity (e.g. through weight loss in mice or symptom reports in humans) and includes immune protection data through challenge studies. Such data would need to be collected for several inoculum doses. Integrated with the models we analyzed here, it could then allow one to predict the impact over a full range of inoculum doses, including those not experimentally measured.

Being able to predict the expected protection achieved for a given inoculum dose can help in the design of vaccines in cases when only limited antigen is available, e.g. in emergency situations [4]. Having information on both immune protection and expected morbidity allows one to determine an optimal inoculum dose based on the - often conflicting - goals of high protection and low morbidity. For instance, one could systematically answer questions such as, “If we require at least 80% immune protection, what would the minimum amount of inoculum need to be? And what level of morbidity/side-effects would this induce?” Currently, both modeling and experiments are not yet able to be used in such a specific manner. However, a tighter integration of experiments with models, and further model refinement should allow one to use the modeling approach discussed here in the future to help design vaccines.

Some promising extensions and refinements of the models are inclusion of further components of the immune response. For instance, given that T-cells are also known to play an important role in immune protection and are affected by inoculum dose [52,53], it would be beneficial to extend experimental and modeling studies in the future and consider both the B-cell and T-cell components of the adaptive response. Similarly, provided more detailed data on specific components of the innate response is available, including those components explicitly in the models might be useful. Another extension would be to consider stochastic models, which would better be able to capture variation among patients. This would require individual host data to be available for analysis and modeling.

To summarize, we developed a modeling framework that might allow a systematic and quantitative determination of the impact of different inoculum doses on resulting immune protection and morbidity. We applied this approach to several data sets to illustrate the general concept and show how it can lead to important insights, e.g. (more inoculum does not always lead to more immune protection’. The modeling and analysis framework presented here can be applied to data from specific vaccine candidates and help to more efficiently determine the optimal dose.

## Acknowledgements

We thank the Antia group members for helpful discussions.

AH, VZ and RA were partially supported by NIH/NIAID grant U19AI117891. KAP was partially supported by a NIH/NIGMS grant P20GM103499, SC INBRE. The funders had no role in study design, data collection and analysis, decision to publish, or preparation of the manuscript.

